# Pump-free and high-throughput generation of monodisperse hydrogel beads by microfluidic step emulsification for dLAMP-on-a-chip

**DOI:** 10.1101/2023.03.12.532292

**Authors:** Jijo Easo George, Riddha Manna, Shomdutta Roy, Savita Kumari, Debjani Paul

## Abstract

Step emulsification (SE), which generates droplets by a sharp change in confinement, has emerged as a potential alternative to flow-focusing technology. Water/dispersed phase is continuously pumped through a shallow inlet channel into a deep chamber pre-filled with the oil/continuous phase. The need for one or more pumps to maintain a continuous flow for droplet generation, and the consequent use of high sample volumes, limit this technique to research labs. Here, we report a pump-free SE technique for rapid and high-throughput generation of monodisperse hydrogel (agarose) beads using <40 *µ*l sample volume. Instead of using syringe pumps, we sequentially pipetted oil and liquid agarose into a microfluidic SE device to generate between 20000 and 80000 agarose beads in ∼ 2 min. We also demonstrated the encapsulation of loop-mediated isothermal amplification mixture inside these beads at the time of their formation. Finally, using these beads as reaction chambers, we amplified nucleic acids from *P. falciparum* and SARS-CoV-2 inside them. The pump-free operation, tiny sample volume, and high-throughput generation of droplets by SE make our technique suitable for point-of-care diagnostics.

## Introduction

The coronavirus pandemic has again highlighted the critical role point-of-care diagnostic tests play in strengthening public health response to disease outbreaks. Besides being sensitive and specific, such tests must also work in low-resource settings and be widely available. Molecular techniques such as RT-qPCR are sensitive, but they require a sophisticated real-time fluorescence thermocycler, which limits their use outside of well-equipped research or pathology labs. Droplet microfluidics allows precise control of the reaction environment with low consumption of reagents.^1,2^ Droplet-based studies with single-cell resolution give better insights into the heterogeneity among cell populations compared to the bulk measurements.^3,4^ Each droplet can act as an independent reaction chamber, leading to high-throughput reactions.

Droplet microfluidics employs co-flow, cross-flow, and flow-focusing modules to generate single or double emulsions.^5–7^ The droplet size and generation frequency depend on fluid properties such as viscosity, interfacial tension, and flow rates. The generation of droplets with a high degree of monodispersity requires pressure controllers and pumps.^8^

Microfluidic step emulsification (MSE), pioneered by Nakajima et al.^9^, has emerged as a viable technique to generate droplets. Droplets form spontaneously when the dispersed phase flows through a long and shallow inlet (i.e., nozzle) into a deep reservoir containing the continuous phase. The term ‘step emulsification’ implies a sudden height change between the inlet microchannel and the reservoir, resembling a step.^10^ This approach is independent of the flow rate in the dripping regime.^11^ It generates droplets based on the geometry of the inlet channel. Depending on the inlet channel shape, these devices can be categorized as the terrace, trapezoid, rectangular, wedge-shaped, and edge-based MSE devices.^9,11–14^

Several reports have focused on improving the monodispersity and throughput of droplet generation. The measures include modifying the channel geometry, parallelizing channels, using a continuous phase with low viscosity, and using electric and magnetic fields.^15–18^ However, reports on microfluidic step emulsification for diagnostic purposes are scarce. Recently, Shi et al. demonstrated digital PCR using vertical step emulsification, which requires a syringe pump, a thermocycler, and a droplet generation set-up that is unsuitable for mass production.^19^ Zhao et al. integrated a step emulsification chip into a PCR tube and generated many monodisperse droplets using continuous flow. However, the complexity of the system requires trained personnel to perform PCR in this device.^20^

We report a portable microfluidic device to generate monodisperse hydrogel (agarose) beads by step emulsification. We used sequential pipetting of continuous and dispersed phases into the chip as an alternative to pumping. Unlike other reported SE devices, the absence of pumps to drive fluid flow minimizes the sample volume required for droplet generation. There are very few reports on droplet or hydrogel bead-based DNA amplification where HFE-7500 engineered fluid is used as the continuous phase. As this fluid is volatile, its evaporation at the amplification temperature destabilizes the droplets or the beads. This problem becomes worse when PDMS, a gas-permeable material, is used to fabricate the droplet generation device. This paper successfully addressed this problem by incorporating a polyethylene (PE) melt into the PDMS chip before amplifying DNA inside the hydrogel beads. We optimized the agarose concentration to increase the throughput of droplet generation. Finally, we performed on-chip isothermal amplification of pathogenic DNA inside the agarose beads to show the suitability of this platform in point-of-care settings.

## Materials and Methods

### Equipment and Chemicals

The photomask with the device design was printed in the nanofabrication facility at the Indian Institute of Science, Bengaluru. A Karl Suss MJB4 mask aligner was used to fabricate the master in the nanofabrication facility of IIT Bombay. 2-inch diameter Si wafers were purchased from Prolyx Microelectronics (Bengaluru, India). A plasma bonding set up comprising of Plasmaflo PDC-FMG-2 and a Plasma Cleaner PDC-32G-2 from Harrick Plasma (USA) was used for bonding the PDMS devices to the glass surfaces. A hot plate (catalog no. 6040) purchased from Tarsons (India) was used as the heating source for isothermal amplification of DNA inside the microfluidic chips. Control isothermal reactions were performed using a Bio-Rad Mini MJ thermocycler. A Nikon Eclipse TiE inverted fluorescence microscope was used to image the droplets. A rheometer (ARES G2 TA/MCR 702) was used to measure the rheological properties of agarose at different temperatures. A JEOL (JSM-7600F) field-emission electron microscope was used to image agarose beads.

SU-8 2005 and SU-8 2010 photoresists were procured from Microchem Corporation (USA). Sylgard 184 polydimethylsiloxane (PDMS) was purchased from Dow Corning Corporation (USA). Novec HFE 7500 engineered fluid from 3M, premixed with an appropriate surfactant, was used as the continuous phase. It was purchased as premixed with 5% FluoSurf (catalog no. 3200811) and with 2% dSurf (catalog no. DR-SE-SU-30) from Dolomite Microfluidics (UK) and Fluigent (France), respectively. 4% (w/w) FluoSurf premixed in HFE 7500 was donated by Emulseo (France) and used for initial optimization experiments. Molecular biology grade water (HiMedia, India) and low-melting agarose (GeNei, Merck) were used as dispersed phases for the generation of droplets and hydrogel beads, respectively. 2019-nCoV_N_Positive Control plasmid (cat. no. 10006625), and the LAMP primers were purchased from Integrated DNA Technologies (USA). WarmStart 2X LAMP kit (DNA and RNA) enzyme mix (catalog no. E1700S) for the loop-mediated isothermal amplification (LAMP) reactions was bought from the New England Biolabs (USA). Unless specified otherwise, all reagents were of analytical grade.

### Device Design

The schematic diagram of the step emulsification device is shown in **figure 1A**. It consists of a single inlet and a single outlet at the opposite ends of a reservoir. The inlet branches into multiple 200 μm long (l) and 20 μm wide (w) inlet channels leading into a 1.25 cm × 2.5 cm reservoir from the two opposite sides. There are 250 inlet channels on either side of the reservoir (shown in the red dashed inset). The height (h) of the inlet channels (shown in the blue dashed inset) is 5 μm, resulting in an aspect ratio (w/h) of 4. The height (H) of the reservoir is 15 μm, resulting in a reservoir-to-channel height (H/h) ratio of 3. There are several circular pillars of 1 mm diameter and a 20 mm × 300 μm rectangular central pillar inside the reservoir to avoid its collapse during plasma bonding owing to the very high width-to-height aspect ratio of the reservoir.

**Figure 1.**
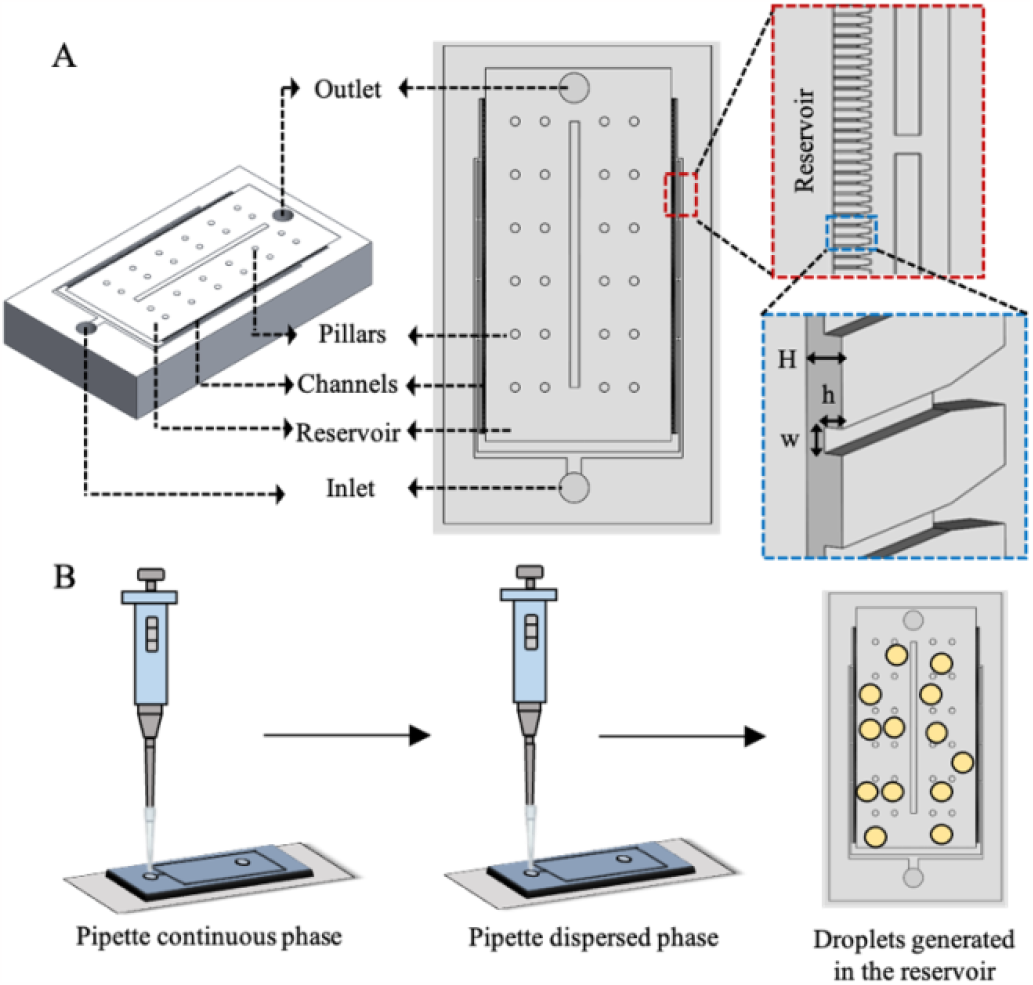
(A) Schematic diagram of the step emulsification device. (B) Schematic diagram showing the generation of droplets by sequential pipetting of continuous and dispersed phases.

### Device Fabrication

The microfluidic device was fabricated by soft lithography. The chrome-on-glass masks were fabricated by laser writing at the Centre for Nano Science and Engineering (CeNSE), Indian Institute of Science, Bengaluru. The SU-8 master was fabricated at the cleanroom of Indian Institute of Technology Bombay’s Nanofabrication Facility by two-layer lithography. A 2-inch Si wafer was first dehydrated at 120°C for 20 min. To pattern the inlet channels (first layer), SU-8 2005 was spin-coated on the wafer in a two-step process: 500 rpm for 15 sec, followed by 2500 rpm for 45 sec. This was followed by soft baking the coated wafer on a hot plate at 95°C for 5 min. The wafer was then exposed to UV (9.88 mW/cm2) for 15 sec through the first mask. Post-exposure bake was done at 95°C for 5 min. After development, the wafer was hard-baked at 95°C for 5 min. For fabricating the reservoir (second layer), the same lithography protocol was used for SU8-2010 and a second mask was used for exposure. PDMS, prepared in the base to curing agent ratio of 10:1, was used to fabricate the device.

In order to minimize evaporation of the continuous phase from the device during DNA amplification, we inserted a 40 μm thick low-density polyethylene (LDPE) barrier sheet into the PDMS layer, following the observations of Ranjith Prakash and others.21 Initially a ∼5 mm thick layer of PDMS was poured on the master and the LDPE film was placed on PDMS. Then the remaining PDMS was poured. Figure S1 in Supplementary Information shows the schematic diagram of the stack containing PDMS and LDPE. The entire stack was cured in a hot air oven at 75°C for 45 min. Then it was heated in the same oven at 135°C for 15 min to melt the LDPE sheet and to allow it to irreversibly adhere to the PDMS surface. The devices were then cut and lifted off from the mold. However, for applications that did not involve DNA amplification, we fabricated PDMS chips without the LDPE insert. Next, we punched inlets/outlets using a 1.5 mm biopsy punch and bonded the devices to a glass cover slip after oxygen plasma treatment for 90 sec. The bonded devices were cured at 65°C for 5 min.

### Generation of water-in-oil droplets and agarose beads by step emulsification

Figure 1B shows the schematic diagram for droplet generation by pipetting. We first pipetted 15 μL of the continuous phase (Novec HFE 7500 premixed with 2% dSurf) into the inlet very slowly until it came out of the outlet, and ensured that it fills the chamber completely, thereby coating the PDMS walls. Subsequently, we pipetted 20 μL of the dispersed phase (i.e., water or agarose solution of a specific concentration) very slowly into the inlet. The presence of a uniform thin film of the continuous phase along the wall ensures that the dispersed phase does not come in contact with the walls of the channel, thus maintaining the radius of curvature of the liquid thread inside the channels and facilitating the generation of monodisperse droplets.

### Characterization of agarose using rheology and FESEM

We used 0.1%, 0.2%, 0.3% and 0.4% (w/v) agarose solutions prepared in distilled water to generate hydrogel beads by step emulsification in our device. We measured the viscosity, the elastic modulus (G’), and the loss modulus (G”) as a function of the shear rate (s-1) and the frequency sweep (Hz) at different concentrations of agarose. Beads were imaged by field emission scanning electron microscopy (FESEM) to measure their pore sizes. Prior to FESEM imaging, the agarose beads were put onto clean coverslips, dehydrated for 1 h and sputter-coated with Pt.

### DNA extraction from *P. falciparum*

RBC cultures containing P. falciparum parasites (0.1% parasitemia) were provided by Prof. Swati Patankar’s group in IIT Bombay. A drop of blood (with and without parasites) was directly spotted on a 3 mm Whatman filter paper and allowed to air dry for storage. DNA was purified from the infected blood spots using Qiagen QIAamp DNA Mini Kit (cat. no. 51304).

### Performing LAMP inside agarose beads

Either the extracted DNA from P. falciparum or the 4012 bp 2019-nCoV_N_Positive Control plasmid, supplied as 2×106 copies/μl, were used as the DNA template for LAMP reactions. LAMP primers designed to target the 18S rRNA region of P. falciparum (a malarial parasite) and N gene of SARS-CoV-2 were used. Six sets of primers were used for each LAMP reaction to increase the efficiency of amplification. These are forward inner primer (FIP), backward inner primer (BIP), forward primer (F3), backward primer (B3), forward loop primer (LF) and backward loop primer (LP). The primer sequences targeting SARS-CoV-2, as reported by Zhang et al22, were as follows:

F3: TGGCTACTACCGAAGAGCT, B3: TGCAGCATTGTTAGCAGGAT,

FIP: TCTGGCCCAGTTCCTAGGTAGTCCAGACGAATTCGTGGTGG,

BIP: AGACGGCATCATATGGGTTGCACGGGTGCCAATGTGATCT,

LF: GGACTGAGATCTTTCATTTTACCGT,

LB: ACTGAGGGAGCCTTGAATACA.

The LAMP primer sets for P. falciparum were as described by Poon et al23:

F3: TGTAATTGGAATGATAGGAATTTA,

B3: GAAAACCTTATTTTGAACAAAGC,

FIP: AGCTGGAATTACCGCGGCTGGGTTCCTAGAGAAACAATTGG,

BIP: TGTTGCAGTTAAAACGTTCGTAGCCCAAACCAGTTTAAATGAAAC,

LF: GCACCAGACTTGCCCT,

LB: TTGAATATTAAAGAA.

A 10X primer mix was prepared with 20 μM of FIP and BIP, 2 μM of F3 and B3, 8 μM of LF and LB. The LAMP reaction mix included 12.5 μl of WarmStart 1X Master mix, 2.5 μl of the 10X primer mix, 1 μl of the proprietary intercalating LAMP Fluorescent Dye and 2 μl of the DNA template. The LAMP Fluorescent Dye can be imaged using the SYBR/FAM channel during fluorescence measurement. The reaction volume was made up to 25 μl by adding nuclease-free water. 5 μl of 1.25% agarose solution was then added to the 25 μl LAMP mix to obtain ∼ 0.2% agarose concentration in the final reaction. The whole reaction was then immediately pipetted into the microfluidic chip that was already filled with oil to generate agarose beads containing the LAMP reaction mixture. On-chip digital LAMP (dLAMP) was performed by directly heating the chip with agarose beads at 75°C on a hot plate for 1 hour. The higher reaction temperature was chosen to compensate for the low thermal conductivity of the glass coverslip. As a positive control, dLAMP was also performed inside the agarose beads by collecting them in a PCR tube and incubating them at 65°C for 1 h in a thermocycler. In both cases, a no-template control (NTC) was used as the negative control for the reaction. The beads were imaged using a Nikon TiE microscope using its standard FITC filter cube (Excitation: 465 - 495 nm, dichroic mirror with a wavelength cut-off at 505 nm, and a barrier filter with a wavelength range of 515 – 555 nm).

## Results and Discussion

### Optimization of device geometry

Droplet generation by microfluidic step emulsification is driven by the surface tension in the dripping regime. When the dispersed phase flows through the narrow microchannel to the edge of the step, the Laplace pressure of the droplet falls rapidly.10,24 This induces a Rayleigh-Plateau instability at the liquid-liquid interface, leading to the pinching of the drop neck.17,25 The dimensionless parameter that determines the regime of droplet generation in a microfluidic device is the capillary number (Ca). If Ca is below a critical value (Ca*), surface tension dominates and droplet generation is independent of the velocity of the dispersed phase (dripping regime).26 In this regime, the size and the monodispersity of the generated droplets are independent of the fluid properties (interfacial tension, viscosity) and the flow velocity.27 It only depends on the aspect ratio ((w/h) > 3) of the inlet microchannel and the radius of curvature of the thread of the dispersed phase inside the channel.28

Initially, we used a transparency mask donated by Neil Davey from the Weitz group (Harvard University).29 Using this mask design, we fabricated a parallel array of 900 µm long, 20 µm wide and 5 µm high wedge-shaped inlet channels that opened into a reservoir of height 50 µm. This device could generate droplets in a high-throughput manner, it led to two problems when generating droplets by pipetting. First, the droplets formed were polydisperse. Second, these droplets were stacked in multiple layers, making their imaging and subsequent analysis difficult.

To improve the monodispersity of the droplets, we ran a parametric simulation in R to explore the dependence of different geometric parameters of the inlet channel and the reservoir on the droplet size. These parameters include length (l), width (w) and height (h) of inlet channel, height of the reservoir (H), and the value of the wedge angle (θ) for wedge-shaped inlet channels. The detailed discussions on how different geometric parameters affect droplet generation by step emulsification are given in the Supplementary Information. The different geometric conditions for generation of spherical and nodoidal droplets are illustrated in figures S2.

We surmised that the observed polydispersity possibly results from incorrect inlet channel dimensions. The volume of the liquid that is added to the growing droplet from the time of the necking of the liquid thread in the inlet channel to the time of its pinch-off to form the droplet (i.e., the excess volume ΔV) increases linearly with the length and the width of the inlet channel, as shown in Figure S3 in the Supplementary Information. The sudden increase in the droplet radius for inlet channel lengths > 300 m and channel widths > 30 m indicates the formation of nodoidal (pancake-shaped) droplets, which we wished to avoid. Since the increase in droplet diameter is due to a dynamic variation in the curvature of the liquid thread in the inlet channel, the inlet channel length is also a critical parameter for optimizing the hydrodynamic resistance. We then optimized the reservoir height (H) to avoid the formation of multiple layers of droplets in the reservoir. We changed the reservoir height such that the condition Δh/h>1 (where Δh = H-h) still holds. The ratio Δh/h determines the aspect ratio of the droplets formed. When the aspect ratio is equal to 1, the droplets are spherical. Otherwise, these droplets are nodoidal30. Theoretically, the aspect ratio w/h of the inlet channel needs to be greater than 3 for droplet generation. Here, we fixed w/h = 4. Considering all the above parameters, we optimized l = 200 m, w = 20 m, and h = 5 m (figures 2A and 2B). Even though a smaller channel length would have sufficed, we chose the channel length to be 200 m to impart sufficient hydrodynamic resistance to the liquid (dispersed phase). As shown in figure S4, the wedge angle needed to be less than 20o for droplet formation. We chose to work with rectangular inlet channels (i.e., with wedge angle 0o) considering fabrication requirements.

**Figure 2.**
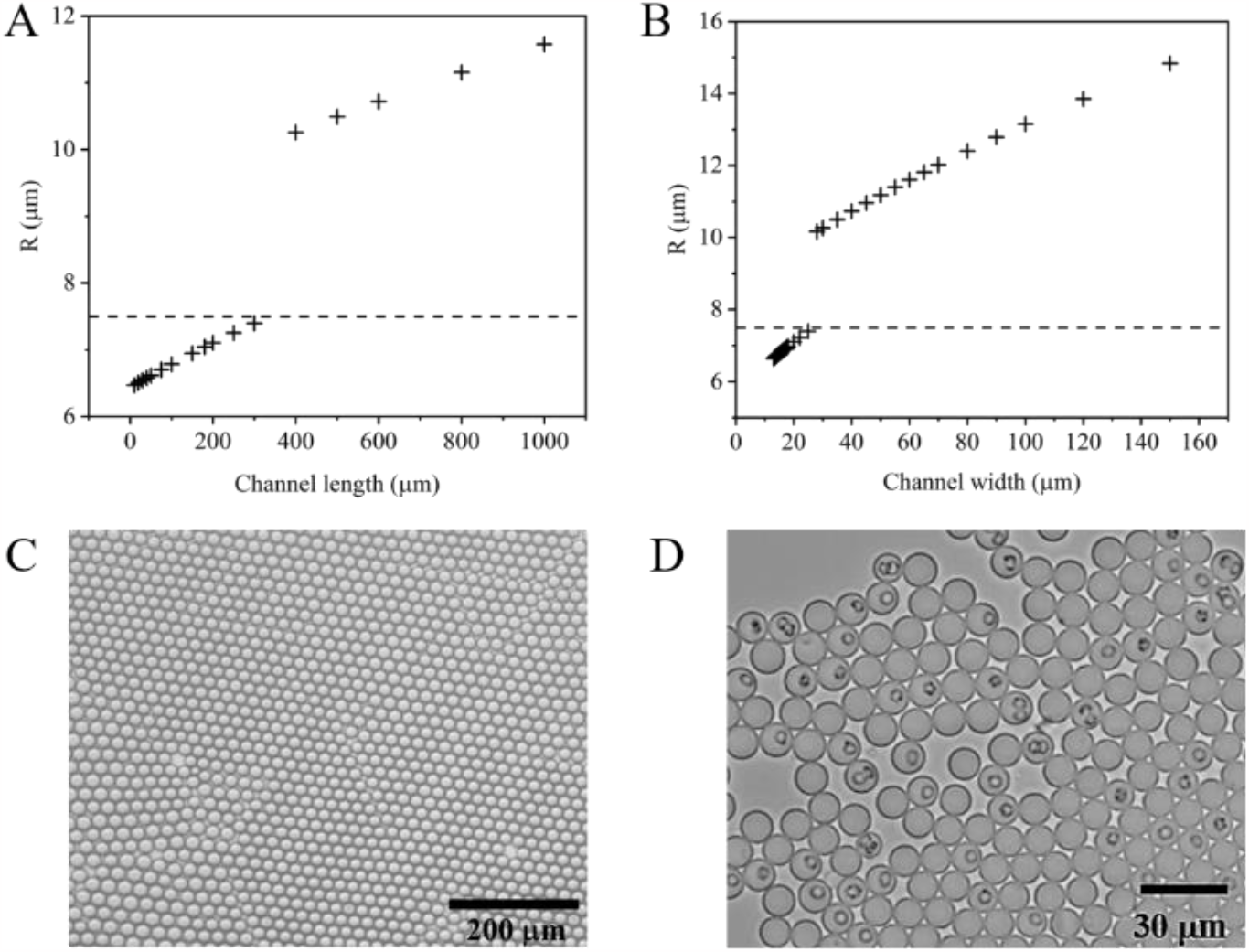
Optimization of the length and the width of the inlet channel. We plotted the variation of the droplet radius as a function of (A) inlet channel length and (B) inlet channel width. The detailed calculations leading to these plots are discussed in the Supplementary Information. (C) Microscope image of water droplets formed in the device fabricated with optimized dimensions. (D) Microscope image of red blood cells (RBS) in phosphate-buffered saline (PBS) droplets.

Theoretically, an appropriate choice of geometrical parameters should ensure that the droplet generation happens in the dripping regime. Experimentally, however, the monodispersity of the generated droplets can also be affected by the large fluctuations in the velocity of the dispersed phase during pipetting. Since monodispersity is theoretically possible only for a specific range of fluid velocity, we also needed to ensure appropriate hydrodynamic resistance in the channel. Table ST1 in Supplementary Information lists the optimized geometrical parameters.

### Generation of water droplets and cell encapsulation

We tested the optimized device design initially by generating water droplets. We first pipetted HFE 7500 mixed with 2% dSurf into the reservoir, followed by slow pipetting of distilled water. We found the average diameter of the water droplets to be ∼ 9 µm (N = 859) with a CV of 8%, indicating a significant improvement in monodispersity due to the improved device design. The experimentally obtained value of droplet diameter is commensurate with the expected value (10 µm) from our parametric simulations. Subsequently, we encapsulated RBCs in the droplets by pipetting 2% whole blood mixed with phosphate buffered saline (PBS) as the dispersed phase. **Figure 2C** shows the microscope image of the water droplets, while **figure 2D** shows single RBCs encapsulated inside the droplets.

### Optimization of agarose concentration for generation of hydrogel beads

Next, we demonstrated pump-free generation of hydrogel (agarose) beads by step emulsification in this device. Hydrogel beads have been used in the past for investigating heterogeneity in proliferation and metastasis at a single-cell level in 3D cell culture as the beads can mimic the extracellular environment.^31,32^ Hydrogel beads can also be useful as chambers for high-throughput biochemical reactions. Further, the reaction products can be stored in the beads until further use. We decided to use agarose beads as reaction chambers because the physical properties of agarose can be tuned easily by varying its concentration.

We first characterized the viscosity (**figure 3A**) and the viscoelastic behaviour (**figure 3B**) of 0.1%, 0.2%, 0.3% and 0.4% agarose gels at 30°C. Figure 3A shows the shear-thinning behaviour of agarose. As expected, the overall viscosity of 0.4% agarose solution was the highest compared to the other concentrations, while 0.2% and 0.3% agarose solutions had similar viscosities for all values of shear rate. As shown in **figure 3B**, we also quantified the elastic modulus (G’) and viscous modulus (G”) of different concentrations of agarose. We noted that the elastic moduli of 0.2% and 0.3% agarose were within a similar range (26 and 38 Pa respectively), as were the loss moduli (10 and 7 Pa respectively), compared with the values for either 0.1% or 0.4% agarose solutions (**table ST2** in **Supplementary Information**). These viscoelastic parameters give an insight into the concentration of agarose gel that can be used in our microfluidic step emulsification device. It is observed that the flow properties of the agarose solution increase concomitantly with the elastic property as the concentration increases from 0.1 - 0.4%. This helps us in concluding that the concentration range used here is appropriate for performing flow experiments in the microfluidics channels without using syringe pumps.

**Figure 2.**
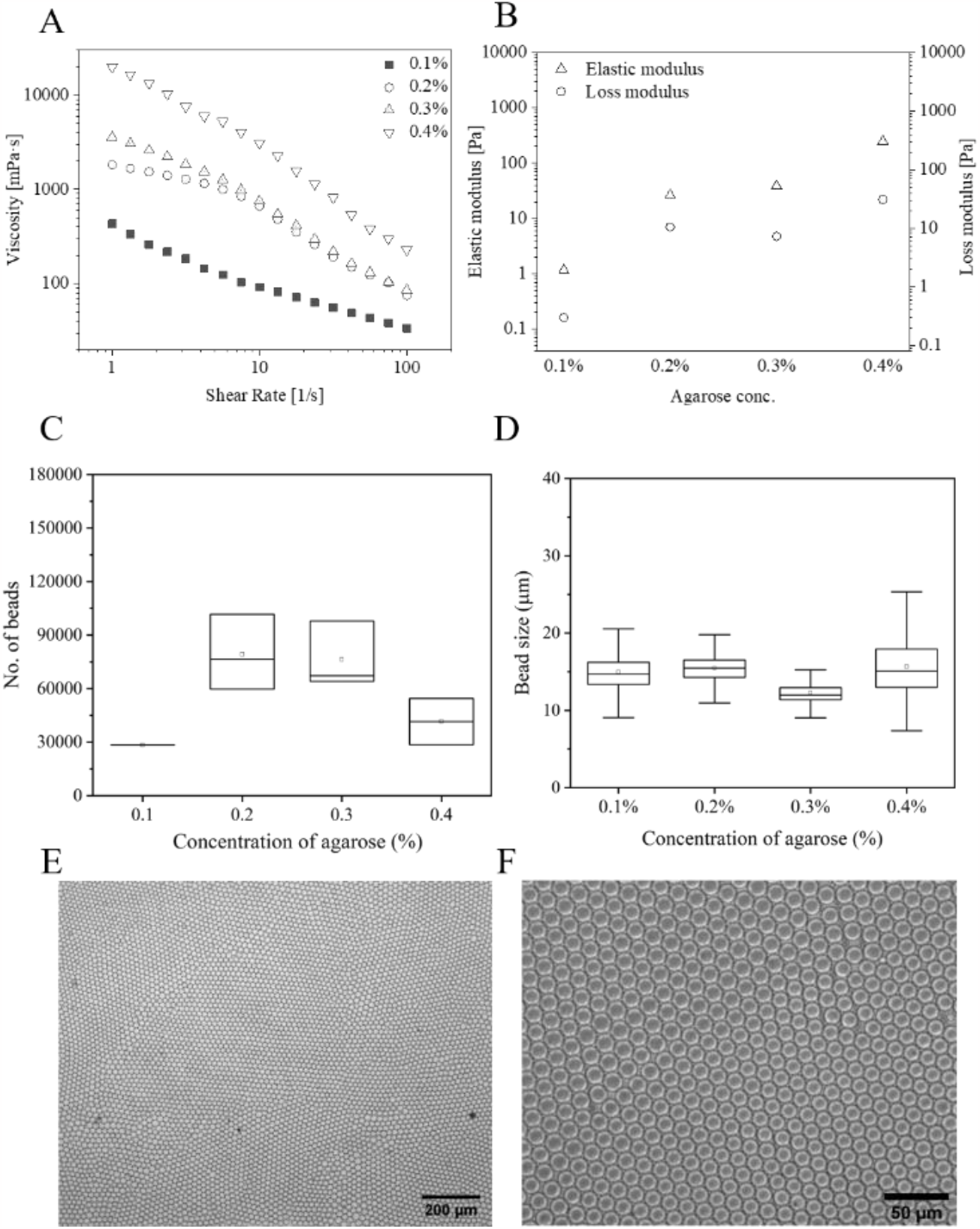
Optimization and characterization of agarose concentration for efficient droplet generation. (A) Variation in the viscosity of agarose as a function of shear rate. (B) Variation of elastic and loss moduli as a function of agarose concentration. Optimization of the agarose concentration in terms of (C) bead number and (D) bead diameter. Microscope images of agarose captured under (A) 4X and (B) 20X objectives.

Subsequently, we generated beads by step emulsification using agarose solutions with concentrations ranging from 0.1% to 0.4%. We found that both 0.2% and 0.3% agarose solutions led to the generation of 75000 - 80000 beads in a single run (figure 3C). For 0.2% agarose, the average number of beads from a single run was ∼ 80,000 at a generation frequency of ∼2666 Hz. The high throughput of our device can be explained by the presence of 500 inlet channels which generate droplets or beads simultaneously. According to the literature, an increase in the viscosity of the dispersed phase results in more inlet channels reaching the breakthrough pressure, thereby increasing the throughput of the device.33 At 0.4% concentration, the agarose solution often solidifies at the inlet, which explains the low throughput at this concentration.

The average diameter (figure 3D) of the agarose beads for all four agarose concentrations is higher than that of the water droplets, ranging from (11.4 ± 0.6) µm (15.6 ± 2.8) µm. This is due to the dependence of the bead diameter on the interfacial tension between the continuous and the dispersed phases. The bead diameter increases with a decrease in the surface tension of the dispersed phase.34 Here, the increase in the average diameter of agarose beads is attributed to the lower interfacial tension of the agarose solution (56 mN/m) compared to that of the DI water (72 mN/m)35. Therefore, 0.2% - 0.3% agarose concentration appears to be the optimum range for generating hydrogel beads using our device. Figures 3E and 3F show the images of the beads captured under 4X and 20X magnifications respectively

### Porosity of agarose beads measured using FESEM

We also quantified the porosity of the agarose beads using their FESEM images. As expected, the average pore size of the agarose beads decreases with an increase in the concentration. The largest average pore size of ∼15 nm corresponded to 0.1% agarose concentration and the smallest average pore size of ∼ 6 nm was obtained for 0.4% agarose concentration (Figure S5 in Supplementary Information). According to the literature, the pore size of 1% agarose gel in the hydrated state is ∼100 nm.36 However, we imaged the agarose beads using a FESEM as we did not have access to a cryo-SEM. The imaging conditions may have shrunk the pores and led to a lower value.

### Amplification of *P. falciparum* and SARS-CoV-2 DNA in agarose beads

Finally, we performed DNA amplification inside the 0.2% agarose beads. Our initial attempts at performing LAMP inside the agarose beads failed due to the following two reasons. First, the beads generated using 2% dSurf coalesced when heated to 75oC for DNA amplification. Second, the engineered fluid evaporated when heated at 75oC for one hour. We made the following changes in our bead generation protocol to address these problems. First, we increased the surfactant concentration from 2% to 5% to avoid coalescence of the beads. Since we did not have dSurf at a higher concentration available with us, we switched to droplet generation using the surfactant FluoSurf for the DNA amplification reactions. As shown in figure 4A, the number of agarose beads generated with 5% FluoSurf was somewhat lower at a value of (55225 ± 2612). The average diameter of the 0.2% agarose beads was (16 ± 2) m when generated using 5% FluoSurf (Figure 4B). Second, we inserted a polyethylene sheet into the PDMS mould as described earlier in the section on device fabrication. Since the implanted PE film was not in direct contact with the reaction mixture, it is not expected to inhibit the amplification.

**Figure 4.**
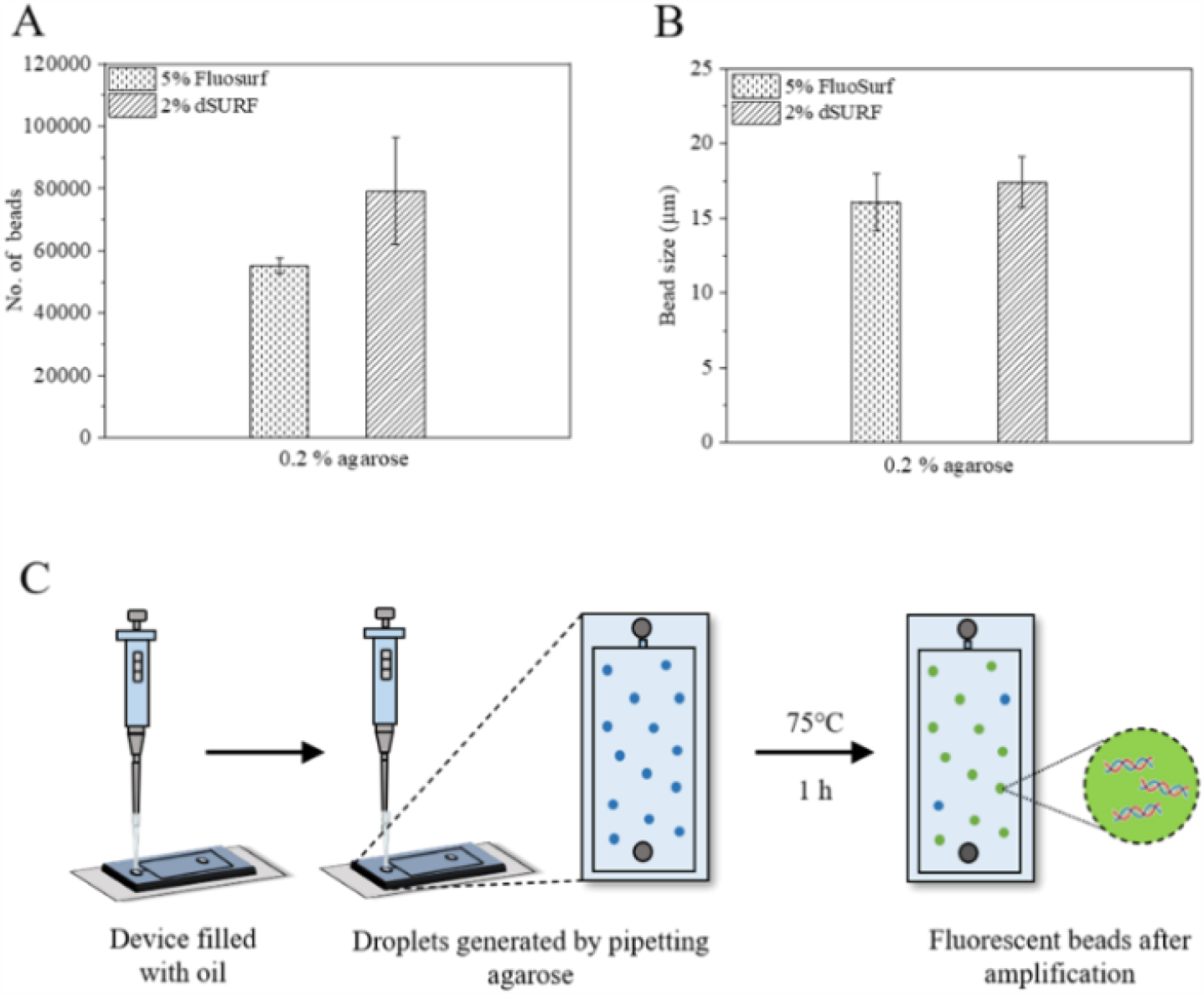
DNA amplification inside agarose beads. (A and B) Comparison of the droplet generation efficiency with surfactants of different concentrations in HFE 7500 oil. (C) Workflow of dLAMP in tube. Fluorescent images of (D) P. falciparum and (E) SARS-CoV-2 after dLAMP in chip. (F) Gel image of amplified DNA.

LAMP is an isothermal amplification technique where the target DNA is amplified at a constant temperature, and hence eliminates the need for a sophisticated thermal cycler. Once the agarose solution with the LAMP reaction mix reaches the step between the inlet channel and the reservoir, it results in the encapsulation of the reaction mixture inside the agarose beads (Figure 4C). The presence of surfactant in the oil stabilizes the thin film formed between the agarose droplets. The use of six primers, specifically the presence of the loop primers, produces several amplicons of different lengths in the gel bead, thereby enhancing the yield of the reaction. Here, we have used DNA from P. falciparum (malarial parasite) and a SARS-CoV-2 (novel coronavirus) plasmid. Successful amplification would lead to the intercalation of the LAMP Fluorescent Dye into amplified dsDNA and would result in fluorescence from the beads. After amplification, the beads were kept at 4°C for 20 min for the sol-gel transition. Then fluorescence images of the agarose beads were captured under a 20X microscope objective (Figures 5A and 5B). We also performed a negative control reaction inside agarose droplets generated in a different chip by omitting the template DNA from the reaction mixture. The presence of fluorescence inside agarose beads depicts the successful DNA amplification of DNA. As expected, the agarose beads in the negative control chip do not show any fluorescence. As additional controls, we performed regular LAMP reactions (i.e., without generating droplets) in the presence of 0.2% agarose inside PCR tubes in a thermocycler. As shown in Figure 5C, the presence of these amplification products was confirmed by running them in a 3% agarose gel. We also observed that the size of the beads decreased in the chip after heating during amplification (Figures 5D and 5E). This observation indicates that there is still partial evaporation in the PDMS chip. Moving to a thermoplastic material would solve this problem completely.

**Figure 5.**
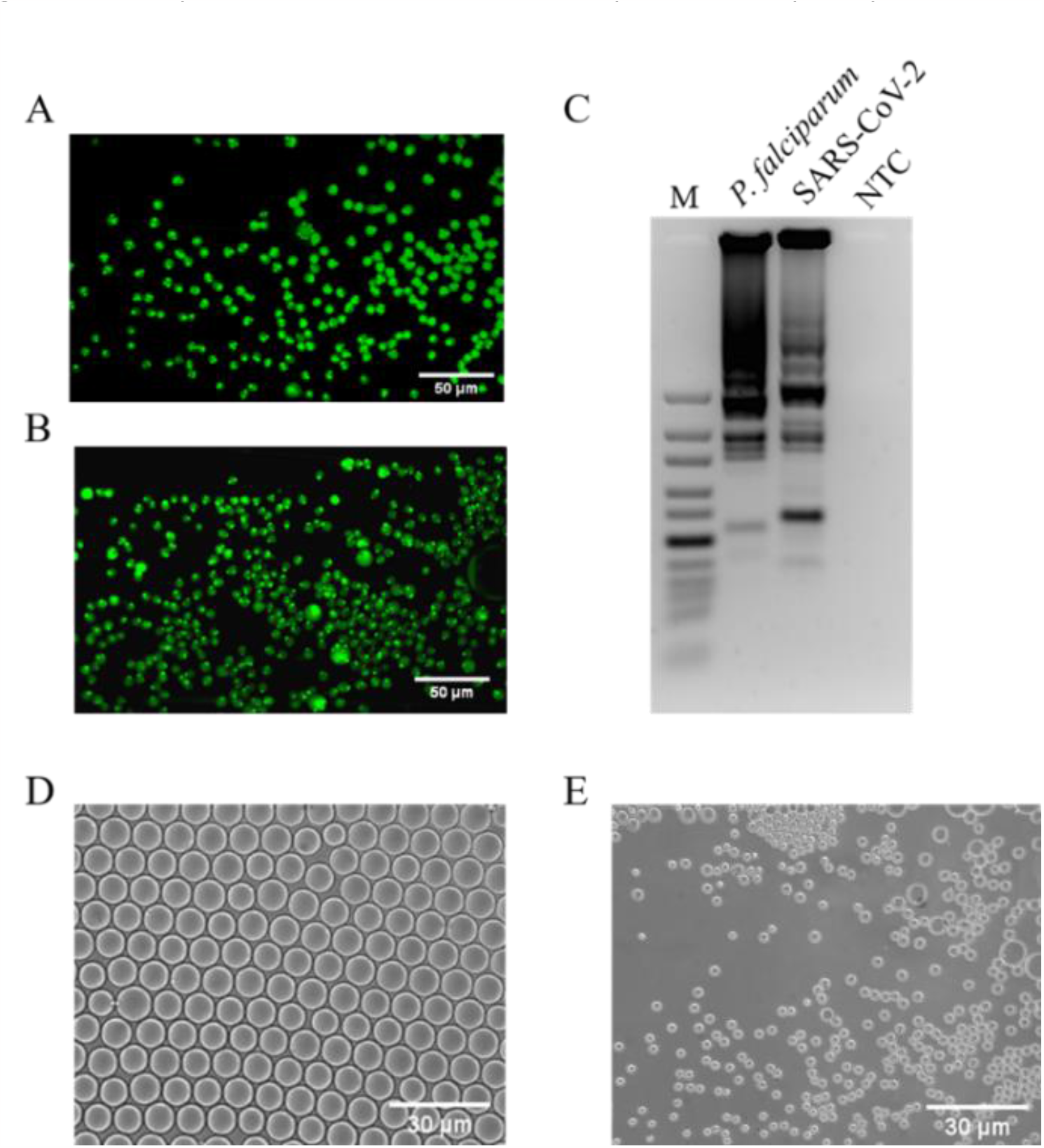
Fluorescent images of (A) P. falciparum and (B) SARS-CoV-2 after DNA amplification on chip. (C) Gel image of amplified DNA. Comparison of droplet size (D) before and (E) after amplification on chip.

We also carried out LAMP reactions by transferring the beads into a closed PCR tube and performing the reaction in a thermocycler instead of a hot plate, as described in the Supplementary Information (Figure S6). This was done to eliminate the effect of evaporation due to heating. The tubes containing beads were also imaged under the fluorescence microscope and indicated successful DNA amplification.

## Conclusions

We report a pump-free technique for high-throughput generation of water-in-oil droplets and agarose beads by microfluidic step emulsification. Our method eliminates the use of syringe pumps and uses a much lower volume of the reagents than what is normally reported. A single run consumes only ∼ 15 l of the expensive engineered fluid and the surfactants. Slow pipetting of the dispersed phase ensures that the droplets are generated in the dripping regime. Our method is much less energy intensive as it does not use pumps. Before fabricating the chip, we optimized its geometrical parameters to improve the throughout and monodispersity of droplets and hydrogel beads. We find that 0.2% - 0.3% agarose concentration range is the most suitable for generation of beads by step emulsification. We also note that the droplet size does not change much over the range of the agarose concentrations that we explored. Finally, we demonstrated the potential of this pump-free technique as a point-of-care (POC) diagnostic tool by amplifying DNA from P. falciparum and SARS-CoV-2 inside these agarose beads. The entire process for droplet generation and DNA amplification involved just two pipetting steps, thereby reducing the chances of manual errors. Most droplet-based microfluidics devices are not used outside of research labs in diagnostic laboratories due to the complex process of droplet generation. Our method paves the way for the development of other POC diagnostic tools based on microfluidic droplet generation.

## Author contributions

Jijo Easo George: Conceptualization, Methodology, Investigation, Formal analysis, Visualization, Validation, Writing – Original Draft; Riddha Manna: Conceptualization, Methodology, Investigation, Formal analysis, Visualization, Writing – Original Draft; Shomdutta Roy: Methodology, Investigation, Writing – Review & Editing; Savita Kumari: Methodology, Investigation; Debjani Paul: Conceptualization, Methodology, Supervision, Funding acquisition, Formal analysis, Writing – Original Draft, Writing – Review & Editing.

## Acknowledgements

JEG acknowledges salary support from an Institute Post-Doctoral Fellowship from IIT Bombay. RM acknowledges a Tata Centre Fellowship from IIT Bombay. The authors acknowledge seed funding from the WHEELS Global Foundation (grant codes DO/2017-SUMP001-001 and RD/0117-DON00G0-001) to start the project, and follow-on funding from the project ‘Infrastructure facility in advanced Research & education in diagnostics’ from the Department of Biotechnology (grant code: RD/0117-DBT0000-013). We acknowledge Priyanka Naik, Sourav Acharya and Anusri U. for technical help. We also gratefully acknowledge Neil Davey and Prof. David Weitz for sharing the initial microfluidic device design. We are grateful to Emulseo for a donation of FluoSurf and to Fluigent for a donation of dSurf, which helped in our initial optimization reactions.

## Supplementary information

### 1. Implanting an LDPE film into PDMS chip

In order to reduce evaporation of the Novec HFE 7500 engineered fluid from the PDMS chip, a LDPE insert was implanted in between the PDMS layers. **Figure S1** shows the schematic diagram of the stack containing the LDPE insert.

**Figure S1.**
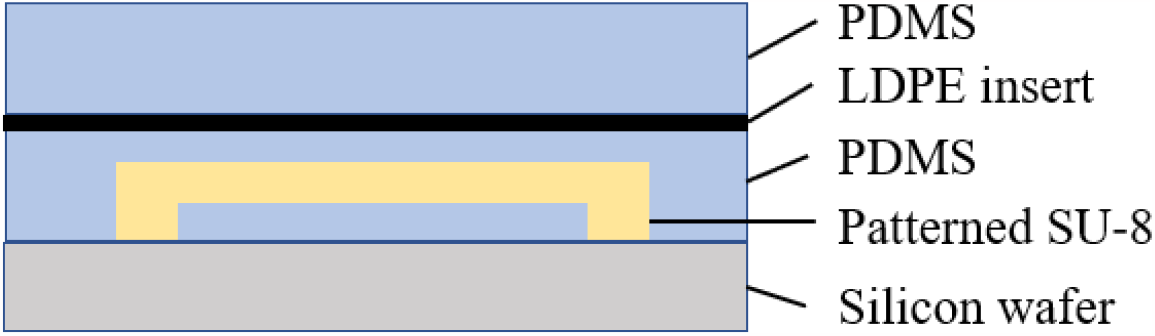
Schematic diagram showing the PDMS stack with the LDPE insert.

### 2. Parameters governing the droplet size

We assume that the dispersed phase is introduced into the reservoir of height **H** through an inlet channel with length ‘**l**’, width ‘**w**’ and height ‘**h**’. The flow rate (**Q**) of the dispersed phase is assumed to be small enough such that droplets are generated in the dripping regime. Let the radius of the confined fluid in the inlet microchannel be **r** and its curvature be **K**_**th**_. The radius of the growing droplet in the reservoir is **R**, while its curvature is **K**_**b**_. The curvatures in all parts of the device must be equilibrated for Laplacian pressure equilibrium. The curvature upstream of the step (i.e., in the inlet channel) is confined by the microchannel geometry. The curvature downstream of the step (i.e., in the reservoir) decreases as the droplet grows in size and becomes less confined. The upstream curvature must adjust to the downstream geometry of the reservoir during droplet formation.

**Figure S2.**
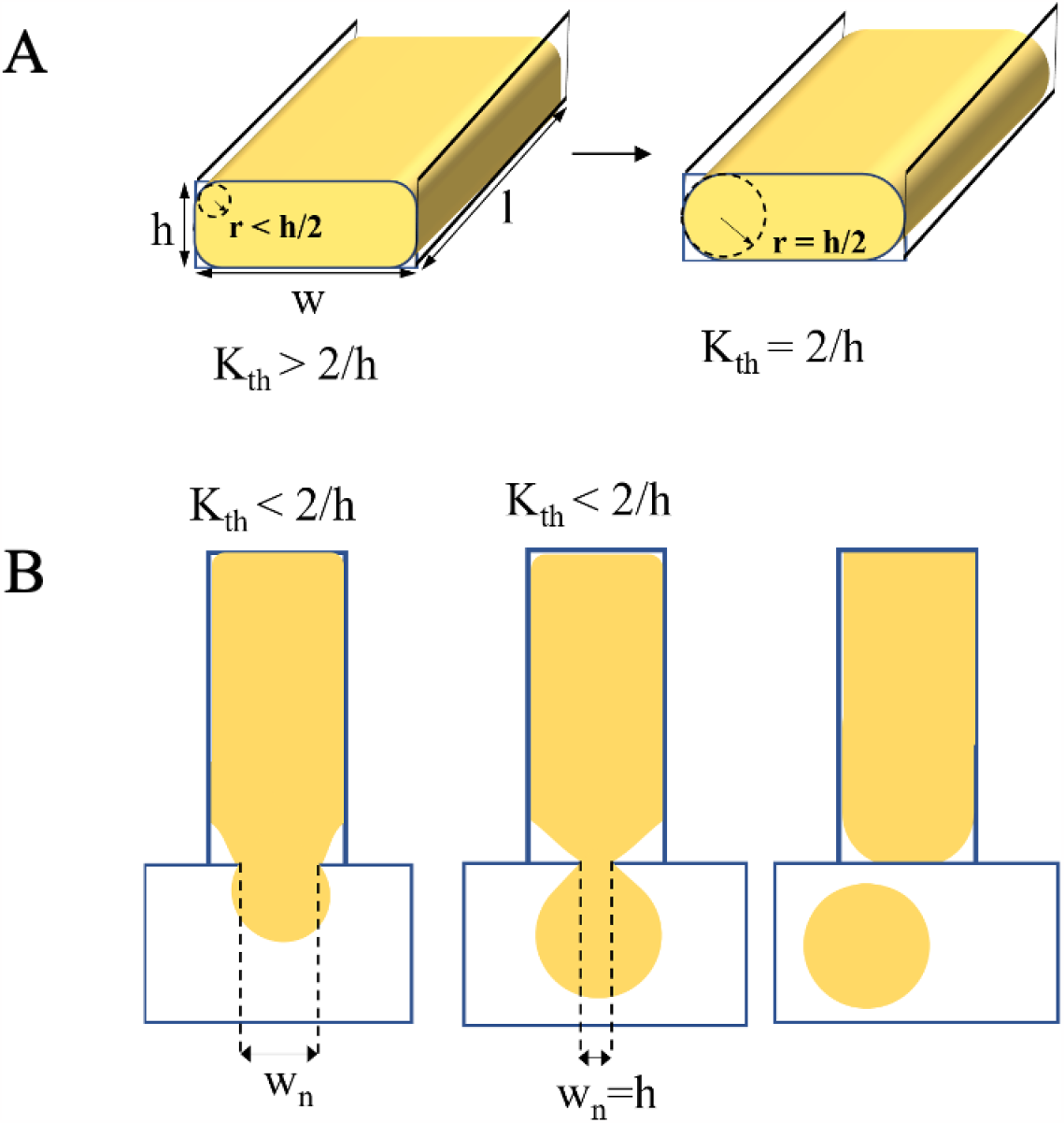
Schematic diagram showing different geometric parameters during droplet generation. (A) Threshold curvature during formation of unconfined and confined droplets. (B) Generation of the liquid bulb and its pinch-off to form the droplet. W_*n*_ indicates the critical width of the neck at which the droplet breaks off from the liquid thread.

The initial radius of curvature (**r**) of the liquid thread in the inlet channel is smaller than the critical radius, **r**_**c**_ **= h/2**. At this time, liquid flows from the inlet channel into the reservoir, generating a bulb. It forms a sphere of radius **R**. As the volume of the liquid in the bulb increases, its radius also increases, and consequently, the curvature, **K**_**b**_ decreases. Mass transport driven by inlet flow happens until the curvature of the bulb (**K**_**b**_) equals the curvature of the liquid thread (**K**_**th**_).

As R increases further, **K**_**b**_ goes down. Due to equilibration of Laplacian pressure, liquid flows into the bulb from the thread, and **K**_**th**_ also keeps decreasing. This happens until both the curvatures reach the critical curvature, **K**_**c**_ **= 2/h**. At this point necking of the thread occurs until the thread becomes cylindrical with **r**_**c**_ **= h/2**. The neck gradually increases in size until the length of the cylinder becomes equal to the critical length, **L**_**c**_ **= *Π*h/2**. At this point, due to Rayleigh-Plateau instability, the droplet pinches off from the thread. The bond number does not play a major role in droplet generation in microfluidic devices, while the capillary number, **Ca** <<1, ensuring quasi-static conditions.

The main geometric parameter governing the size of the droplets is the critical curvature (**K**_**c**_) of the bulb which depends on **h**, the height of the inlet channel. The height of the reservoir (**H**) also plays a vital role in determining the shape, and hence the radius (**R**), of the droplet. We define **Δh = H – h**. If **Δh/h > 1**, spherical droplets are formed. **H** also determines the aspect ratio (***χ* = 2R/H**) of the droplets. If **Δh/h ≥ 1, *χ* = 1**, indicating spherical droplets.

The length of the inlet channel (**l**) influences the extra volume (ΔV_1_) added during transition of **K**_**th**_ from its initial value to the critical value (**K**_**c)**_.

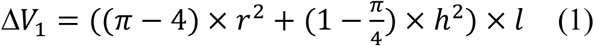

The width (**w**) of the inlet channels influences the extra volume (ΔV_&_) that is added during the necking of the thread.

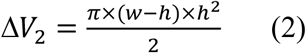

Under unconfined conditions, the droplet remains a sphere. For spherical droplets, the total volume (**V**) is given by equation (3).

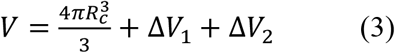

The radius (**R**) of the droplet is given by equation (4).

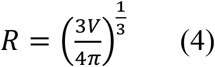

For nodoidal droplets, the volume (**V**) is given by equation (5).

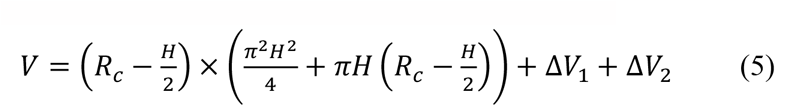

The radius (**R**) of nodoidal droplets is given by equation (6).

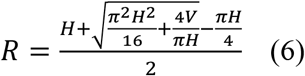

Using the above expressions, a script was written in R to compare the changes in the droplet volume (ΔV) and the droplet radius (ΔR) due to variations in inlet channel length (l) and inlet channel width (w). The corresponding graphs were plotted in **figure S2**. For all the simulations, the height of the inlet channel (**h**) and the height of the reservoir (**H**) were kept constant at 3 *µ*m and 4.5 *µ*m, respectively. When the channel length was varied, the width was kept constant at 12 *µ*m. When the width was varied, the channel length was kept constant at 100 *µ*m.

The funnel-shaped step was incorporated in the first-generation device design to make the device passive, i.e., to make the droplet radius (**R**) independent of the flow rate of the dispersed phase. A funnel-shaped inlet also helps in the back flow of the continuous phase, thus aiding in the pinching off of the droplet from the fluid inside the microchannel. A simple geometrical analysis shows the effect of the wedge angle on the pinching off of the droplet. **Figure S3** shows two situations for a spherical droplet of a fluid with a contact angle ***ϕ***, placed in a wedge of wedge angle ***θ***. In **figure S3(A)**, the droplet does not wet the apex of the wedge when **2*ϕ* > (180° + *θ*)**. In **figure S3(B)**, it wets the apex of the wedge when (**180° - *θ*) < 2*ϕ* < (180° + *θ*)**.

**Figure S3.**
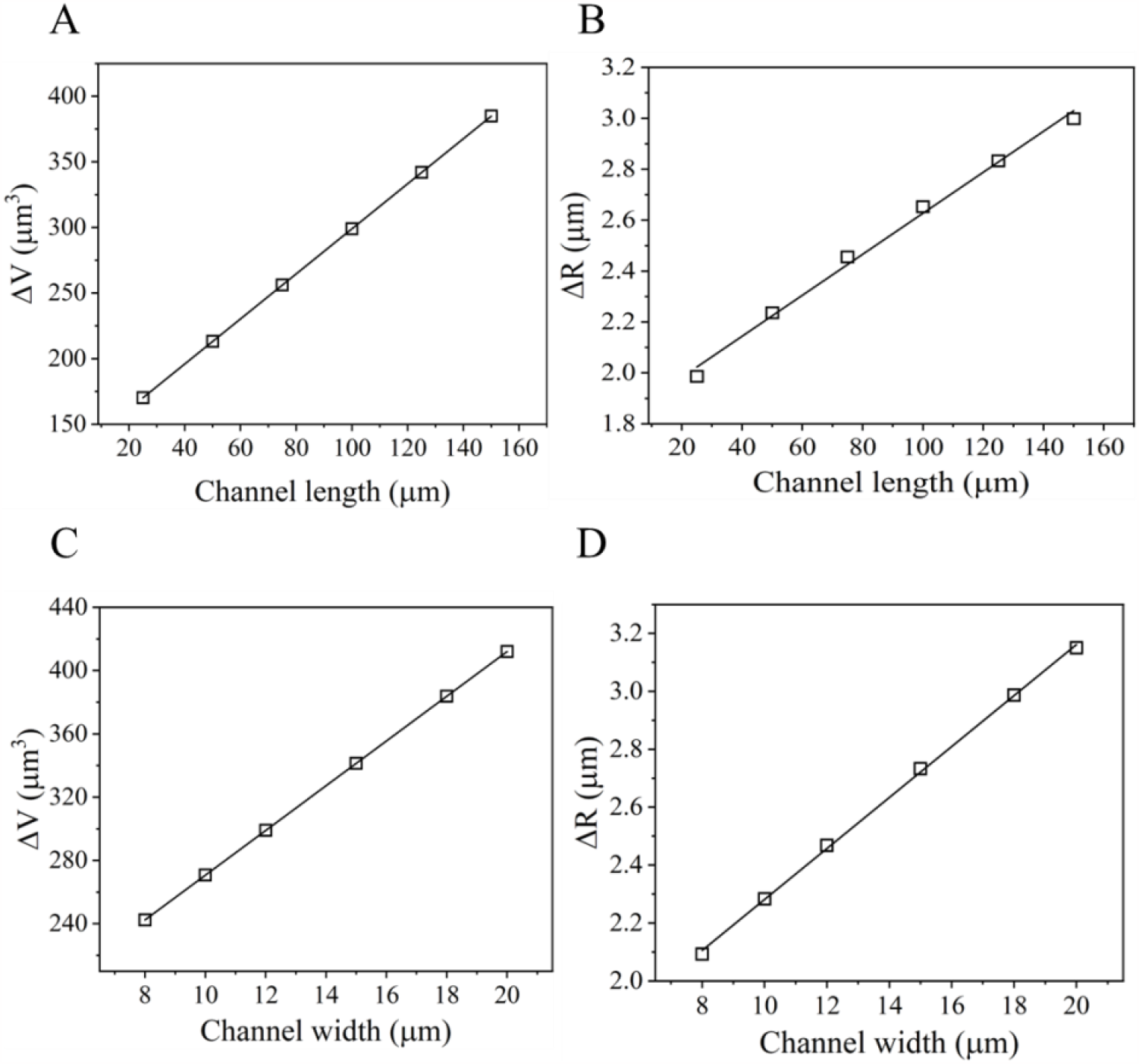
(A) Change in the excess volume as a function of channel length. (B) Change in droplet radius as a function of channel length. (C) Change in the excess volume as a function of channel width. (D) Change in droplet radius as a function of channel width.

**Figure S4.**
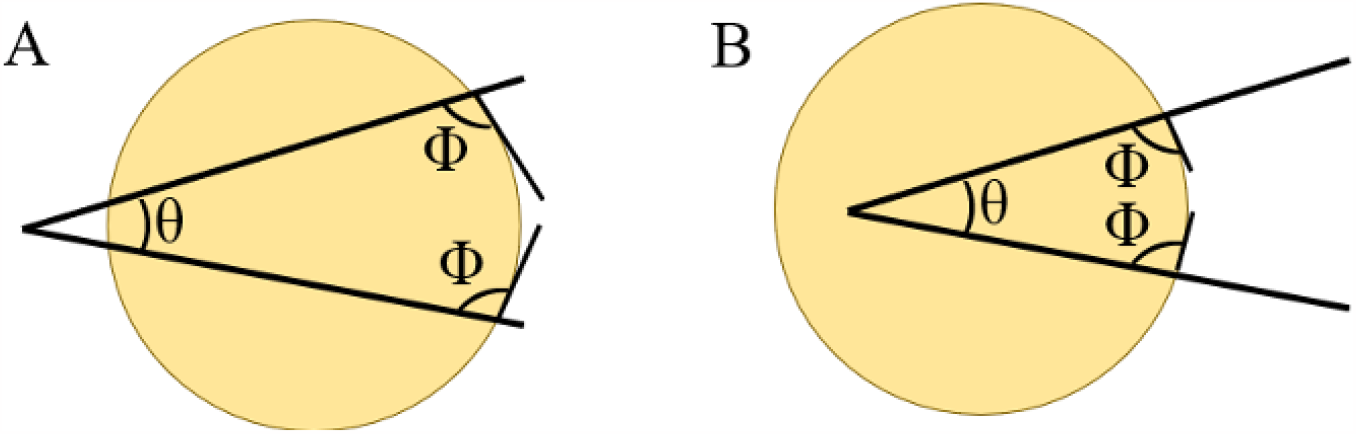
(A)The droplet does not wet the apex of the wedge when 2ϕ > 180° + θ. (B) The droplet wets the apex of the wedge when 180° - θ < 2ϕ < 180° + θ.

For the device geometry to aid in droplet formation, it should not favour the wetting of the apex of the wedge by the diluted blood sample. Hence, we much have

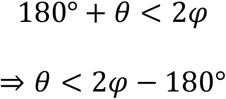

We found that diluted blood (5%) has a contact angle of ∼100° with PDMS surface. Plugging in this value in the above equation, we get θ < 20° for the funnel-shaped geometry to favour droplet formation. We decided to go with a rectangular inlet channel with wedge angle 0° as fabricating the wedges accurately was difficult. **Table ST1** summarizes the various optimized geometrical parameters.

**Table ST1:** Optimized geometrical parameters for the device

| Parameter | Initial values<br>(first-generation<br>device) | Final values (our<br>simulation) |
| --- | --- | --- |
| Length of inlet<br>channel (l) | 900 $\mu\text{m}$ | 200 $\mu\text{m}$ |
| Width of inlet<br>channel (w) | 20 $\mu\text{m}$ | 20 $\mu\text{m}$ |
| Height of the inlet<br>channel (h) | 5 $\mu\text{m}$ | 5 $\mu\text{m}$ |
| Reservoir height (H) | 15 $\mu\text{m}$ | 50 $\mu\text{m}$ |
| Wedge angle | $< 20^\circ$ | $0^\circ$ |

### 3. Optimization of the agarose concentration

The elastic and the loss moduli of different agarose concentrations were measured in order to decide the most suitable agarose gel concentration to be used for the generation of droplets.

**Table ST2.** Elastic and loss moduli of different concentrations of agarose gel.

| Agarose<br>conc. (%) | Elastic<br>modulus (Pa) | Loss<br>modulus (Pa) |
| --- | --- | --- |
| 0.1 | 1.15 | 0.3 |
| 0.2 | 26.31 | 10.5 |
| 0.3 | 38.6 | 7.3 |
| 0.4 | 248.6 | 30.8 |

### 4. Porosity measurement of agarose beads using FESEM

The porosity of the agarose beads was measured from the FESEM images as shown in **figure S4**.

**Figure S5.**
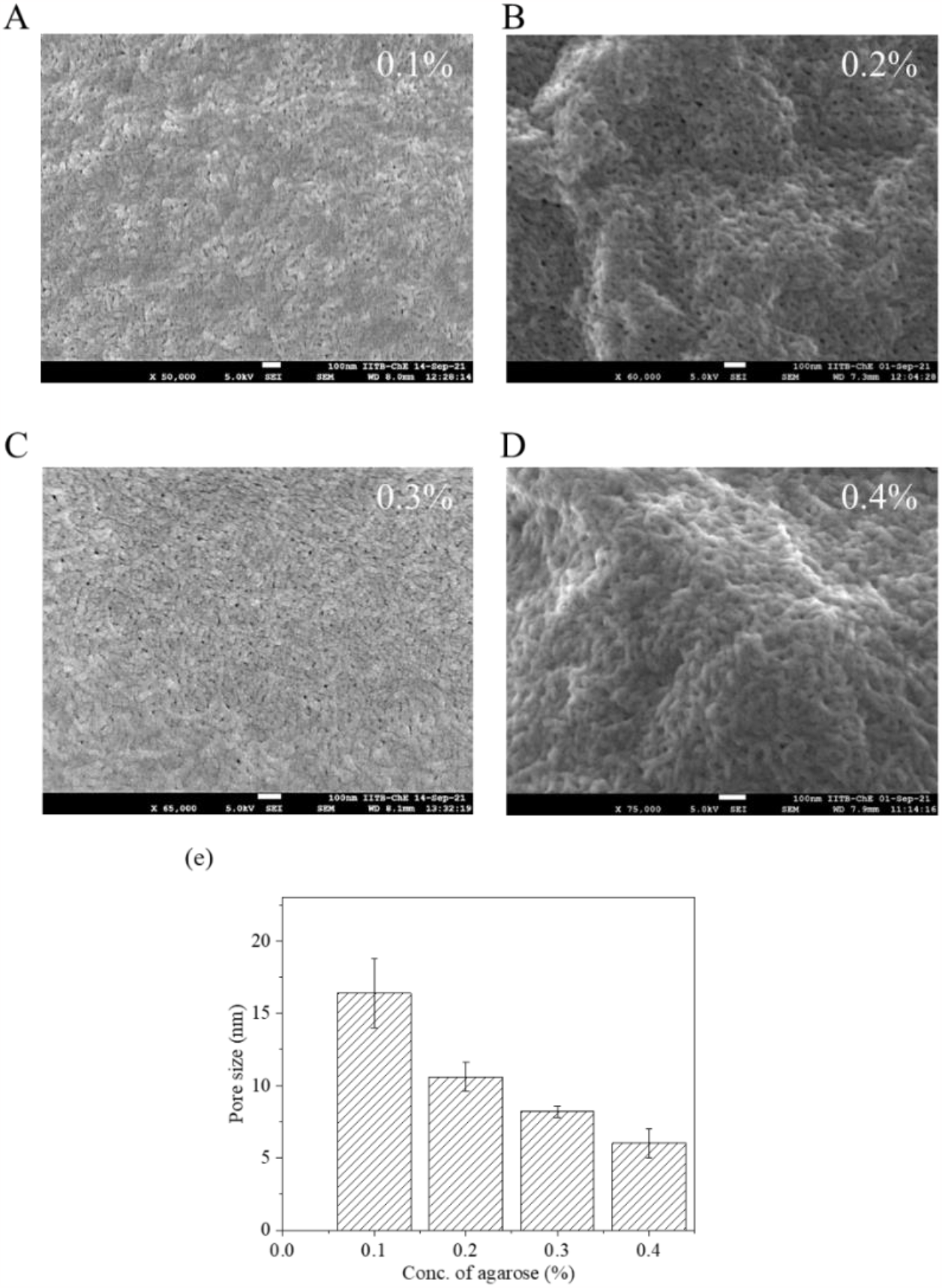
Determining pore size of agarose beads at different concentrations. FESEM images of (A) 0.1%, (B) 0.2%, (C) 0.3%, (D) 0.4% agarose beads. (C) Plot showing the decrease in pore size with increase in agarose concentration.

### 5. DNA amplification in agarose beads inside a closed PCR tube

Agarose beads, containing the amplification reaction mix and the proprietary fluorescent dye present in the LAMP kit, were generated in the microfluidic chip. These beads were then pushed out of the device by pipetting excess oil from the inlet and collected in a PCR tube. DNA from *P. falciparum* and a SARS-CoV-2 plasmid were amplified by LAMP at 65°C for 1 h in a thermal cycler, followed by cooling at 4°C for 20 min. **Figure S5A** shows the schematic diagram of the workflow. The tubes were then directly placed under a fluorescent microscope and the beads were imaged using the FITC filter (**Figures S5B** and **S5C**). Presence of fluorescent beads in the image indicated successful amplification. However, this method requires additional user intervention of transferring the gel beads into the tube, which may lead to contamination.

**Figure S6.**
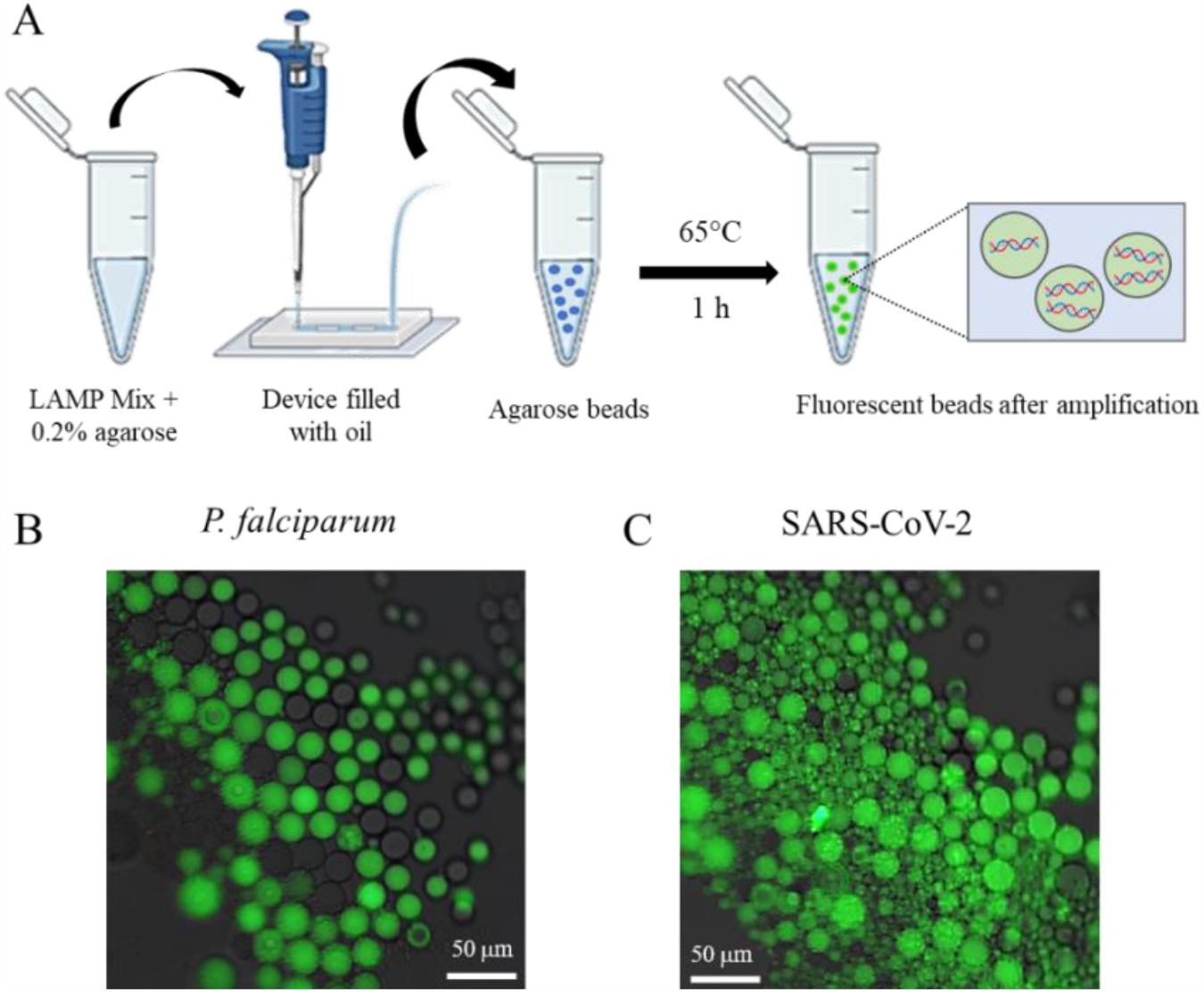
DNA amplification in agarose beads generated in the device. (A) Workflow of dLAMP in tube. Fluorescent images of (B) P. falciparum and (C) SARS-CoV-2 after dLAMP in tube.

